# Hyd ubiquitinates the NF-κB co-factor Akirin to activate an effective immune response in *Drosophila*

**DOI:** 10.1101/323170

**Authors:** Alexandre Cammarata-Mouchtouris, Xuan-Hung Nguyen, François Bonnay, Akira Goto, Amir Orian, Marie-Odile Fauvarque, Michael Boutros, Jean-Marc Reichhart, Nicolas Matt

## Abstract

Upon microbial infection in *Drosophila*, the E3-ubiquitin ligase Hyd ubiquitinylates the NF-κB co-factor Akirin for its efficient binding to the NF-κB factor Relish and subsequent activation of immune effectors genes.

**ABSTRACT:** The *Drosophila* IMD pathway is activated upon microbial challenge with Gramnegative bacteria to trigger the innate immune response. In order to decipher this NF-κB signaling pathway, we undertook an *ex-vivo* RNAi screen targeting specifically E3 ubiquitin ligases and identified the HECT E3 ubiquitin ligase Hyperplastic Discs “Hyd” as a new actor of the IMD pathway. We showed that Hyd targets the NF-κB cofactor of Akirin. The K63-polyubiquitination chains deposited by Hyd decorate Akirin for its efficient binding to the NF-κB transcription factor Relish. We showed that this Hyd-mediated interaction is critical to activate immune-induced genes that depend on both Relish and Akirin, but is dispensable for those that depend solely on Relish. Therefore Hyd is key in operating a NF-κB transcriptional selectivity downstream of the IMD pathway. *Drosophila* depleted for Hyd or Akirin failed to express the full set of immune-induced anti-microbial peptide coding genes and succumbed to immune challenges. We showed further that Ubr5, the mammalian homolog of Hyd, is also required downstream of the NF-κB pathway for the IL1β-mediated *IL6* activation. This study links the action of a E3-ubiquitin ligase to the activation of immune effector genes, deepening our understanding of the involvement of ubiquitination in inflammation and identifying a potential target for the control of inflammatory diseases.

## INTRODUCTION

During evolution, metazoans developed strategies to effectively protect themselves from microbial threats. The similarity between the molecular pathways mediating the innate immune response in insects and mammals points to *Drosophila* as a relevant model to explore the immune response (*1, 2*). In *Drosophila*, the defense against microbes is ensured mainly by the massive production of antimicrobial peptides (AMP) (*3*). Their expression is under the control of two transcription factors belonging to the NF-κB family: Dorsal-related Immunity Factor (DIF) and Relish, acting downstream of Toll and IMD pathways respectively. They are the homologues of mammalian RelB and p50 transcription factors.

Post-translational regulation of proteins by ubiquitin pathway is key for proper immune response, albeit not fully understood (*4*). Conjugation of ubiquitin polymers to proteins by an ubiquitin-ligase is a key mechanism for controlling their activity or stability. Lysine (Lys) residues of proteins can be modified by a polymer of ubiquitin (polyubiquitin) linked through Lys 48 (K48) or Lys63 (K63) of the ubiquitin molecule. Whereas K48-linked polyubiquitin mainly triggers degradation of proteins by the proteasome, K63-linked polyubiquitin regulates, mainly through modification of interactions, the activity and the subcellular localization of proteins (*5*). In both mammals and *Drosophila*, ubiquitination is involved at various levels of the NF-κB pathways (*6*). Furthermore, deregulation of ubiquitin-ligases is implicated in inflammatory pathologies (*7, 8*) and tumor progression (*9*).

In *Drosophila*, IAP2 is the only E3-ubiquitin ligase identified so far as a positive regulator of the IMD pathway (*10, 11*). This protein is involved in the formation and activity of upstream protein complexes formed around the IMD protein and the IKK kinase. To deepen our understanding of NF-κB pathway regulation by the ubiquitination system, we focused on identifying *Drosophila* new ubiquitin-ligases required for the activity of the IMD pathway through a RNAi-based screen in *Drosophila* S2 cells.

Several E3-ubiquitin ligases emerged from this screen as positive or negative regulators of the IMD pathway. We decided to focus on Hyd as i) it is the unique HECT E3-ubiquitin ligase potentially involved in the IMD pathway and ii) it has also emerged as a potential IMD pathway regulator in a parallel pilot screen undertaken in our laboratory (unpublished data from Dr. Akira Goto and (*12*). Our data showed that Hyd is required *in-vivo* to survive an immune challenge with Gram-negative bacteria. Epistasic analysis revealed that Hyd acts at the level of the NF-κB co-factor Akirin, which is known to orchestrate the activation of a subset of NF-κB target genes in combination with the SWI/SNF chromatin remodeling complex (*13-15*). This is consistent with the described localization of Hyd within the nucleus (*16, 17*).

We showed that Hyd decorates Akirin with K63-polyUb chain, which is required for Akirin binding to the NF-κB factor Relish. Furthermore, we observed that Ubr5 (also known as EDD1), the human ortholog of Hyd (*18*), has a conserved function in NF-κB signaling in human HeLa cell line. Similarly to human-Akirin2, Ubr5 is required for the activation of only a subset of NF-κB target genes. We demonstrate here that upon immune challenge, ubiquitin chains are instrumental to bridge NF-κB and its co-factor Akirin to activate an effective immune response.

## RESULTS

### Hyd is an **E3**-ubiquitin ligase required for the activation of the IMD pathway

To uncover novel E3-ubiquitin ligases that modulate the IMD pathway, we screened a library of 174 double strand RNA (dsRNA) targeting putative E3-ubiquitin ligases encoded in the *Drosophila* genome as described in Flybase (*19*). We used stably transfected *Drosophila* S2 cells expressing the *Attacin-A-luciferase* gene, a reporter of the activation of the IMD pathway upon immune challenge with Gram-negative bacteria (*20*). We evaluated the ability of dsRNA targeting individually each of the 174 putative E3 ubiquitin-ligases to interfere with the IMD reporter upon stimulation by heat-killed *Escherichia coli* (HKE), a regular IMD pathway agonist.

*IAP2* is an E3-ubiquitin ligase that positively regulates the pathway by targeting IMD and DREDD (*21*). The knockdown of *IAP2* resulted in a strong decrease of the *Attacin-A-luciferase* reporter induction upon immune stimulation regarding to *dsGFP* control (Fig 1A), providing proof of concept for the screen. Knockdown of six E3-ubiquitin ligase-coding genes (*m-cup, Mkrn1, CG2926, CG31807, mura* and *CG12200*) resulted in a strong increase in *Attacin-A-luciferase* activity upon immune stimulation. Therefore these E3-ubiquitin ligases behave as negative regulators of the IMD pathway. Conversely, the knockdown of three genes encoding either two Really Interesting New Gene (RING) domain E3-ubiquitin ligases *bon* and *CG5334*, or HECT domain E3-ubiquitin ligase *hyd*, resulted in a significant decrease of *Attacin-A-luciferase* activity (Fig 1A). This suggests that Bon, CG5334 and Hyd are new positive regulators of the IMD pathway. We decided to focus on the exploration of Hyd, as it is the unique HECT domain E3-ubiquitin ligase involved in the *Drosophila* IMD pathway.

**Figure 1.**
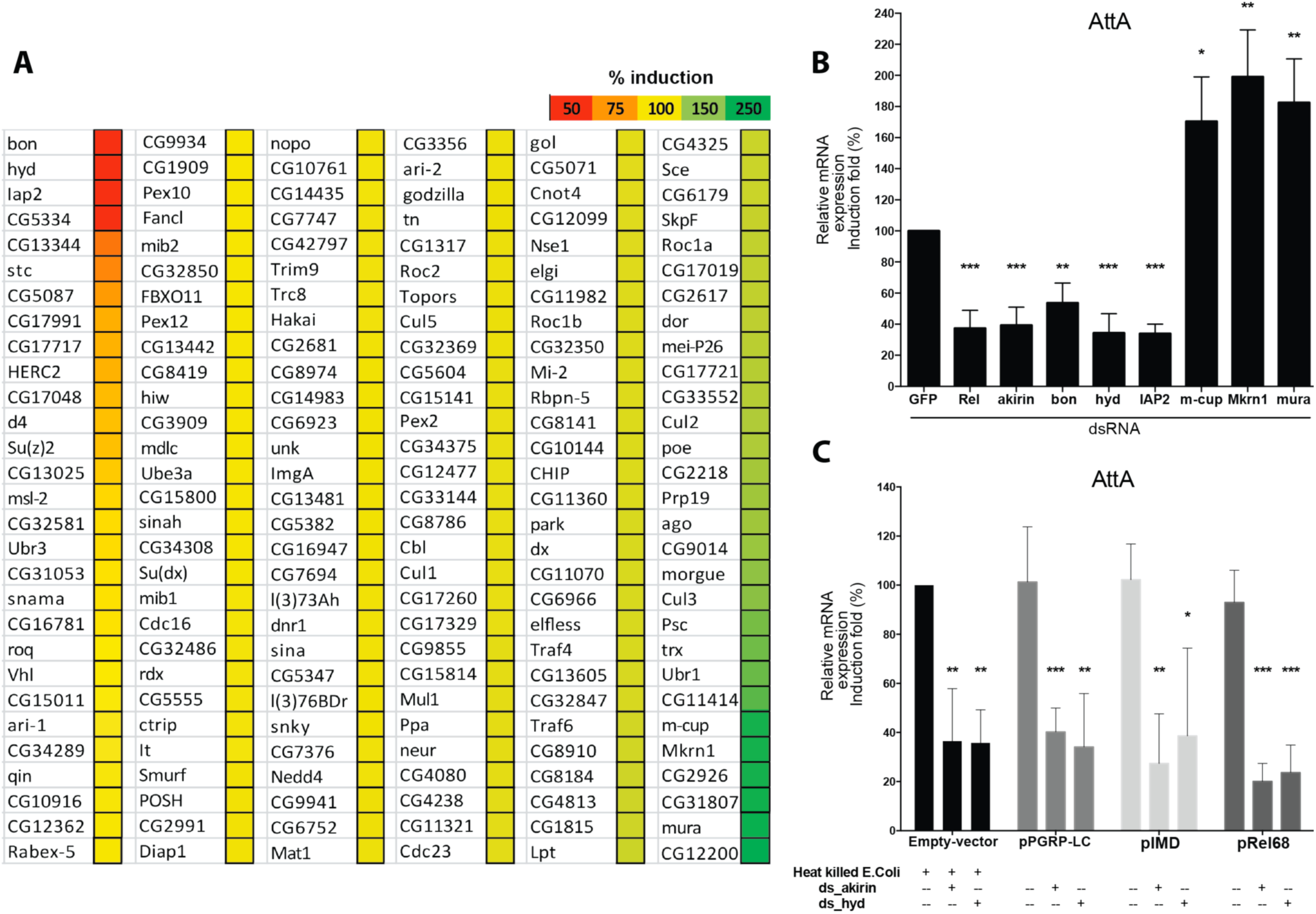
E3-ubiquitin ligases screen identified *ex-vivo* Hyd as involved in IMD pathway. (A) E3 ubiquitin-ligases screen in *Drosophila* S2 cells realized by luciferase assay. The different genes were knocked down by dsRNA. Induction of IMD pathway was done by 48h HKE stimulation and assessed by measure of *Attacin-A* and put on percentage compared to control (dsGFP). (B) Quantitative RT-PCR of *Attacin-A* mRNA from S2 cells transfected with dsRNA against GFP (negative control), relish, akirin (positive controls) and some E3-ubiquitin ligases from the screen, following 4h of HKE stimulation. (C) Epistasis analysis of Hyd position within the IMD pathway. The IMD pathway was induced by either HKE stimulation or the transfection of S2 cells with PGRP-LC, IMD-V5 or Rel-HA plasmids. Cells treated with vector alone serve as a control. Cells were also transfected with dsRNA targeting akirin or hyd. Data information: Data are represented as mean ± standard deviation of three independent experiments realized on 5 × 10^5^ cells (B-C) per sample. Statistical significance was established by comparing values from stimulated with unstimulated conditions and genes knockdown with GFP dsRNA control. *P-value < 0.05; **P-value < 0.01; ***P-value < 0.001.

To validate reporter-assay experiments, *Drosophila* S2 cells were transfected with *dsRNA* targeting either the NF-κB factor *relish*, its cofactor *akirin, hyd* or some of the other E3-ubiquitin ligases of the screen. We challenged S2 cells with HKE and monitored endogenous *Attacin-A* mRNA level by RT-qPCR. Interfering with *relish, akirin* or *hyd* expression significantly decreased HKE-mediated *Attacin-A* induction, compared to control (*dsGFP*) (Fig 1B, Fig S1). We observed that the RING-domain E3-ubiquitin ligases Bon, CG5334, m-cup, Mkm1 and Mura are required for the normal activation of *Attacin-A* expression and that the HECT E3-ubiquitin ligase Hyd acts as a positive regulator of *Attacin-A* expression in *Drosophila* S2 cells (Fig 1B).

In order to identify at which level of the IMD pathway Hyd is required, we undertook an epistasis analysis. *Drosophila* S2 cells were treated by dsRNA targeting *hyd* or *akirin* as a control and the IMD pathway was activated at different levels by transfecting either a truncated form of PeptidoGlycan Receptor Protein-Long Chain a (PGRP-LCa), IMD or the 68kD active-form of Relish (Rel68) (*13*). Measurement of *Attacin-A* expression by RT-qPCR assessed activation of the IMD pathway. We could show that Hyd is required at the same level or downstream of Relish (Fig 1C) to exert its positive regulation on IMD pathway activation.

### Hyd acts at the level of Akirin to trigger full activation of the IMD pathway

Downstream of the IMD pathway, Relish target genes are divided in two subsets: genes that depend only on Relish for their expression (including *Attacin-D* and the majority of negative regulators) and ones requiring Akirin in addition to Relish (including *Attacin-A* and the majority of effectors) (Bonnay, Nguyen et al., 2014). Upon immune challenge in S2 cells, using RT-qPCR, we observed that Hyd depletion recapitulates the immune phenotype of cells depleted for Akirin (Fig 2A). Consequently, Hyd is acting on Akirin-dependent NF-κB transcriptional selectivity *ex-vivo*.

**Figure 2.**
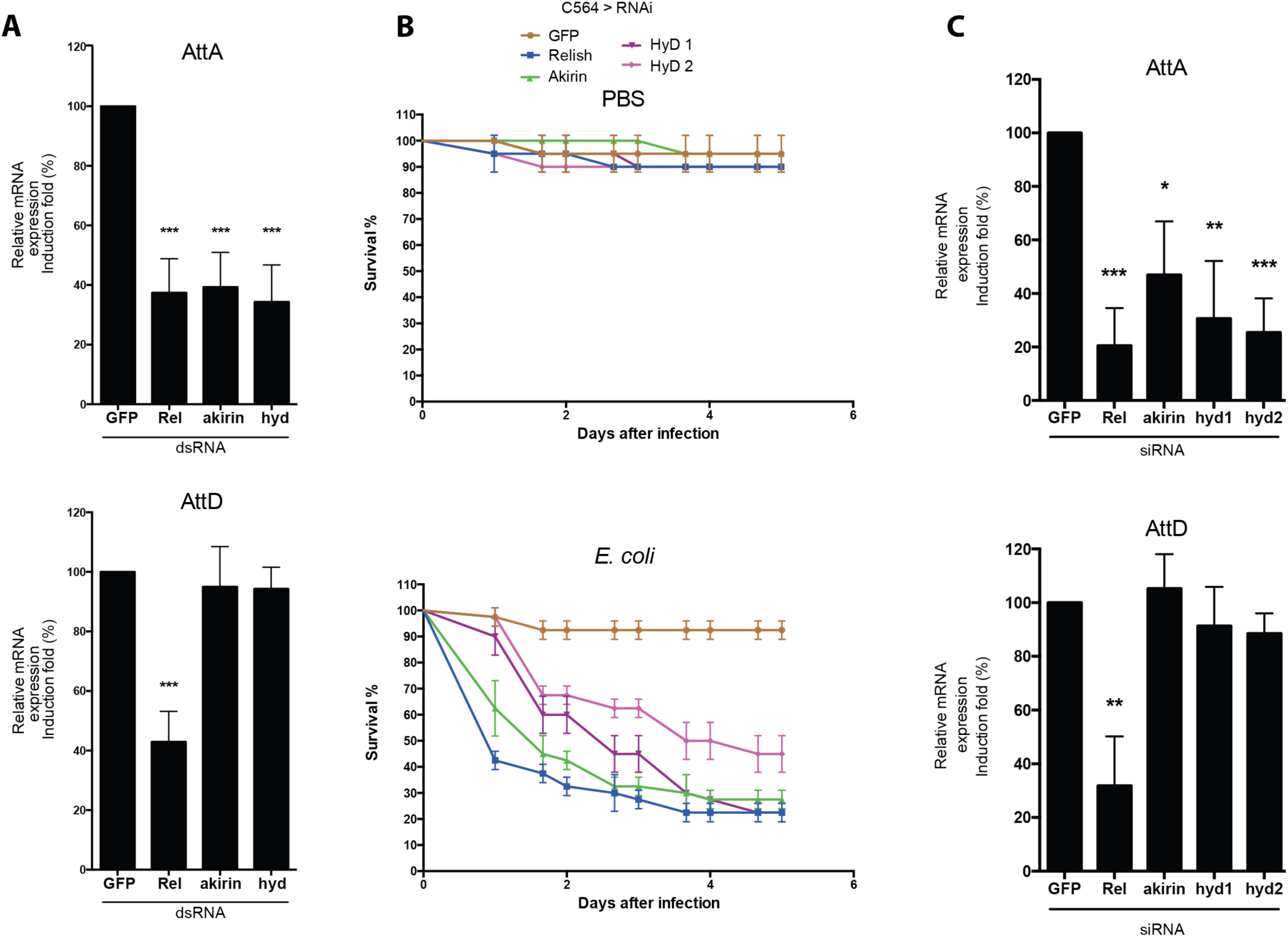
Hyd is required for the full activation of IMD response. (A) Quantitative RT-PCR of *Attacin-A* and *Attacin-D* mRNA from S2 cells transfected with dsRNA against GFP (negative control), relish, akirin (positive controls), and hyd, following 4h of HKE stimulation. (B) *In-vivo* survival experiments performed on batches of 20 nine-day-old females infected by E. coli septic injury (with PBS pricking as control), at 25°C three independent times. Quantitative RT-PCR of *Attacin-A* and *Attacin-D* mRNA, from three batches of 10 nine-day-old males infected with E. coli for 6h by septic injury at 25°C, three times independently. Data information: Data are represented as mean ± standard deviation of three independent experiments realized on 5 ×10^5^ cells (A and C) per sample. Statistical significance was established by comparing values from stimulated with unstimulated conditions and genes knockdown with GFP dsRNA control. *P-value < 0.05; **P-value < 0.01; ***P-value < 0.001.

We next investigated if Akirin and Hyd were similarly required for NF-κB transcriptional selectivity *in-vivo*. As *Drosophila* embryonic development is impaired in absence of Akirin, we used the *C564-Gal4* transgene (*22*) to express RNAi constructs targeting *akirin, hyd* and *relish* in the adult fat body, the main immune organ of *Drosophila* (*3*). Flies depleted of Akirin (*C564 > RNAi-akirin*), Relish (*C564 > RNAi-relish*) or Hyd (*C564 > RNAi-hyd1* or *C564 > RNAi-hyd2*) displayed an impaired survival following *E. coli* infections when compared to control flies (*C564 > RNAi-GFP*) or following PBS pricking (Fig 2B).

Following immune challenge by *E. coli*, expression of *Attacin-A*, but not of *Attacin-D*, was reduced in the absence of Akirin or Hyd, when compared to control flies (*C564 > RNAi-GFP*) (Fig 2C, Fig S2).

Our results indicate that Hyd is required at the level of Relish to activate the Akirin-dependent subset of Relish target genes during the immune response, allowing *Drosophila* to survive a Gram-negative bacterial challenge.

### Hyd mediated K63-polyubiquitination of Akirin is instrumental for its link to Relish

We next investigated if Akirin could be a *bona-fide* target for the E3 ubiquitin-ligase Hyd. Co-immunoprecipitation assay in S2 cells showed that V5-tagged Hyd (*Hyd-V5*) (*23*) binds to endogenous Akirin (Fig 3A). By contrast V5-tagged HydCS (*HydCS-V5*), which displays a mutated HECT domain by conversion of the catalytic cysteine at position 2854 to serine (*23*), is unable to bind to Akirin (Fig 3A). As a control we confirmed that IAP2, the E3-ubiquitin ligase acting upstream of Akirin in the IMD signaling cascade (*10, 11*), does not interact with Akirin (Fig S3).

**Figure 3.**
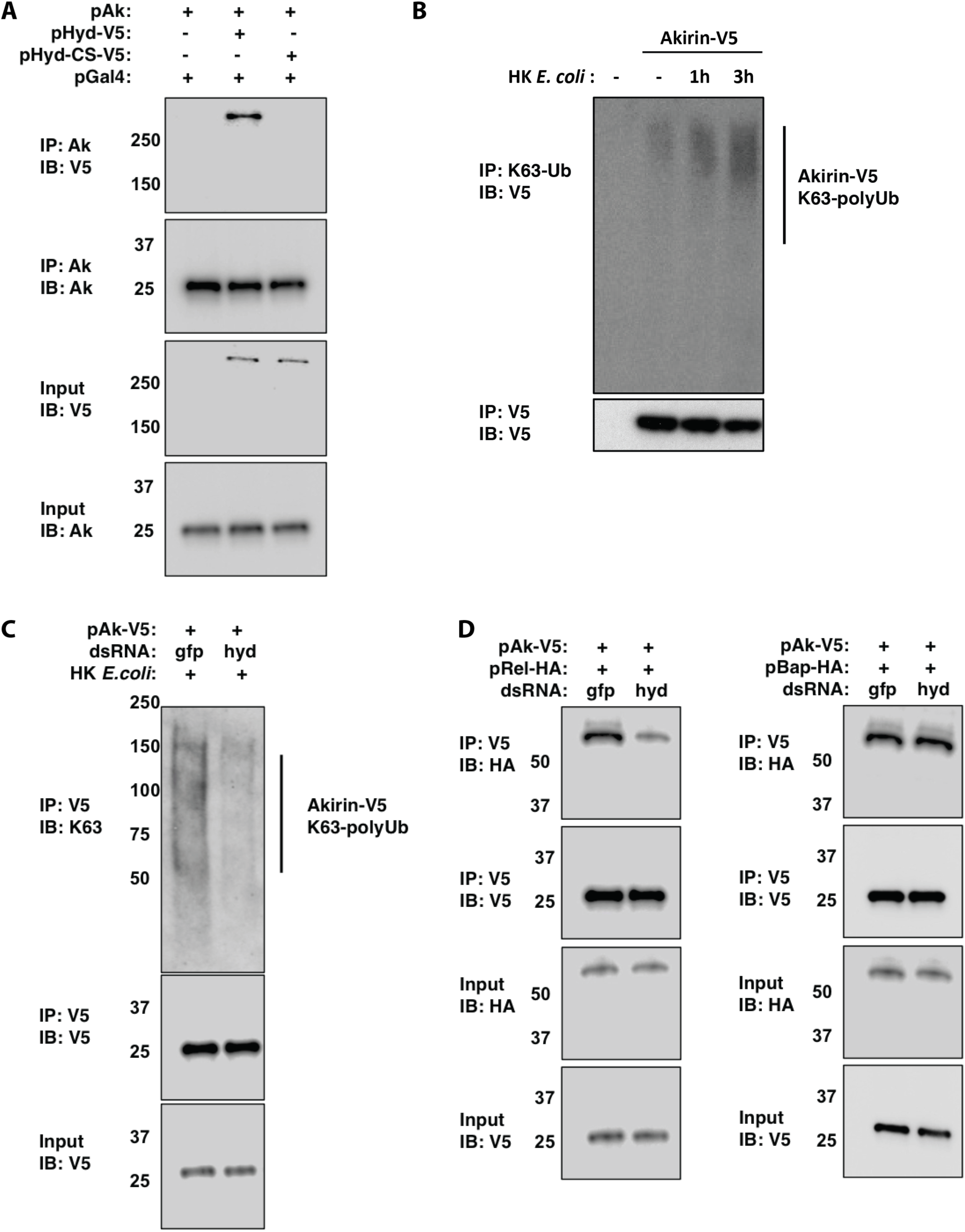
Hyd mediated-ubiquitination of Akirin is necessary for interaction with Relish. (A) Co-immunoprecipitation assay between over-expressed Akirin and Hyd in S2 cells. The cells were transiently transfected with pAC-Akirin, pGal4 and/or pUAS-Hyd-V5 and pUAS-Hyd-CS-V5. Cell lysates were immunoprecipitated with anti-Akirin coupled agarose beads. Immunoprecipitates were analyzed by Western blotting with anti-V5 or anti-Akirin antibodies. (B) Immunoprecipitation assay of K63-polyUb chains on Akirin before and after immune challenge (1h and 3h HKE). S2 cells were transiently transfected with pAC-Akirin-V5. Cell lysates were immunoprecipitated with anti-K63-polyUb coupled agarose beads. Immunoprecipitates were analyzed by Western blotting with anti-V5 antibodies. (C) Immunoprecipitation assay of Akirin after immune challenge (4h HKE). S2 cells were transiently transfected with pAC-Akirin-V5 and dsRNA targeting GFP or hyd. Cell lysates were immunoprecipitated with anti-V5 coupled agarose beads. Immunoprecipitates were analyzed by Western blotting with anti-K63-polyUb and anti-V5 antibodies. (D) Co-immunoprecipitation assays between over-expressed Akirin and Relish or Bap in S2 cells. The cells were transiently transfected with pAC-Akirin-V5 and pMT-Rel-HA or pMT-Bap-HA; and dsRNA targeting GFP or hyd. Cell lysates were immunoprecipitated with anti-V5 coupled agarose beads. Immunoprecipitates were analyzed by Western blotting with anti-HA or anti-V5 antibodies. Data information: Data are representative of 2 independent experiments.

Protein extracts from cells transfected with a tagged version of Akirin (*Akirin-V5*) were immunoprecipitated with an anti-V5 antibody. Western-blot experiments with antibodies targeting K63-polyUb chains suggested that Akirin is K63-polyubiquitinalyted 1h and 3h after immune challenge with HKE (Fig 3B). This immune-induced post-translational modification of Akirin is indeed attenuated upon knockdown of Hyd (Fig 3C). Collectively, these data indicate that upon immune challenge, Hyd physically interacts with Akirin through its catalytic HECT domain to decorate Akirin with K63-polyUb chains. We previously published that Akirin physically bridges the NF-κB factor Relish and BAP60, a core member of the SWI/SNF chromatin-remodeling complex (*14*). To understand whether Akirin K63-polyubiquitination is instrumental for the interaction of Akirin with Relish or BAP60, we performed co-immunoprecipitation experiments in S2 cells depleted for Hyd and transfected with *Akirin-V5* and *Rel68-HA* or *BAP60-HA* (Fig 3D). As previously reported (*14*), Akirin-V5 co-precipitated either with the active form of the NF-κB factor Relish (Rel68-HA) or with BAP60 (BAP60-HA) (Fig 3D). However, in the absence of Hyd, the interaction between Akirin-V5 and Rel68-HA is weakened (Fig 3D). Of note the interaction between Akirin-V5 and BAP60-HA is independent of Hyd (Fig 3D). These results indicate that Hyd is required to deposit K63-polyUb chains on Akirin for subsequent binding to the NF-κB factor Relish.

### Ubr5 - the human ortholog of Hyd - is required for the NF-κB transcriptional selectivity during the inflammatory response

The Akirin-dependent molecular mechanism underlying the selective activation of NF-κB target genes is well conserved from *Drosophila* to mammals (*13-15*). Therefore, we addressed the potential requirement of Ubr5 (the ortholog of the *Drosophila* E3-ubiquitin ligase Hyd) in NF-κB selective transcriptional response mediated by hAkirin2 during the human inflammatory response. We depleted HeLa cells for either NF-κB 1, hAkirin2 or Ubr5 by siRNA (using scrambled siRNA as controls). We monitored, upon stimulation by IL1β, the expression levels of NF-κB target genes that are dependent of hAkirin2 (such as *IL6*) or independent (such as *IL8*) (*15*). As expected (*15*), lacking NFkB1 in HeLa cells impaired both *IL6* and *IL8* activation upon IL1β stimulation. However, the activation of *IL6* and *IL8* is uncoupled in HeLa cells depleted for hAK2 or Ubr5 (Fig 4A, Fig S4). This result suggests a conserved function of Ubr5 in the selective transcription of NF-κB target genes mediated by hAkirin2 that remains to be functionally explored.

**Figure 4.**
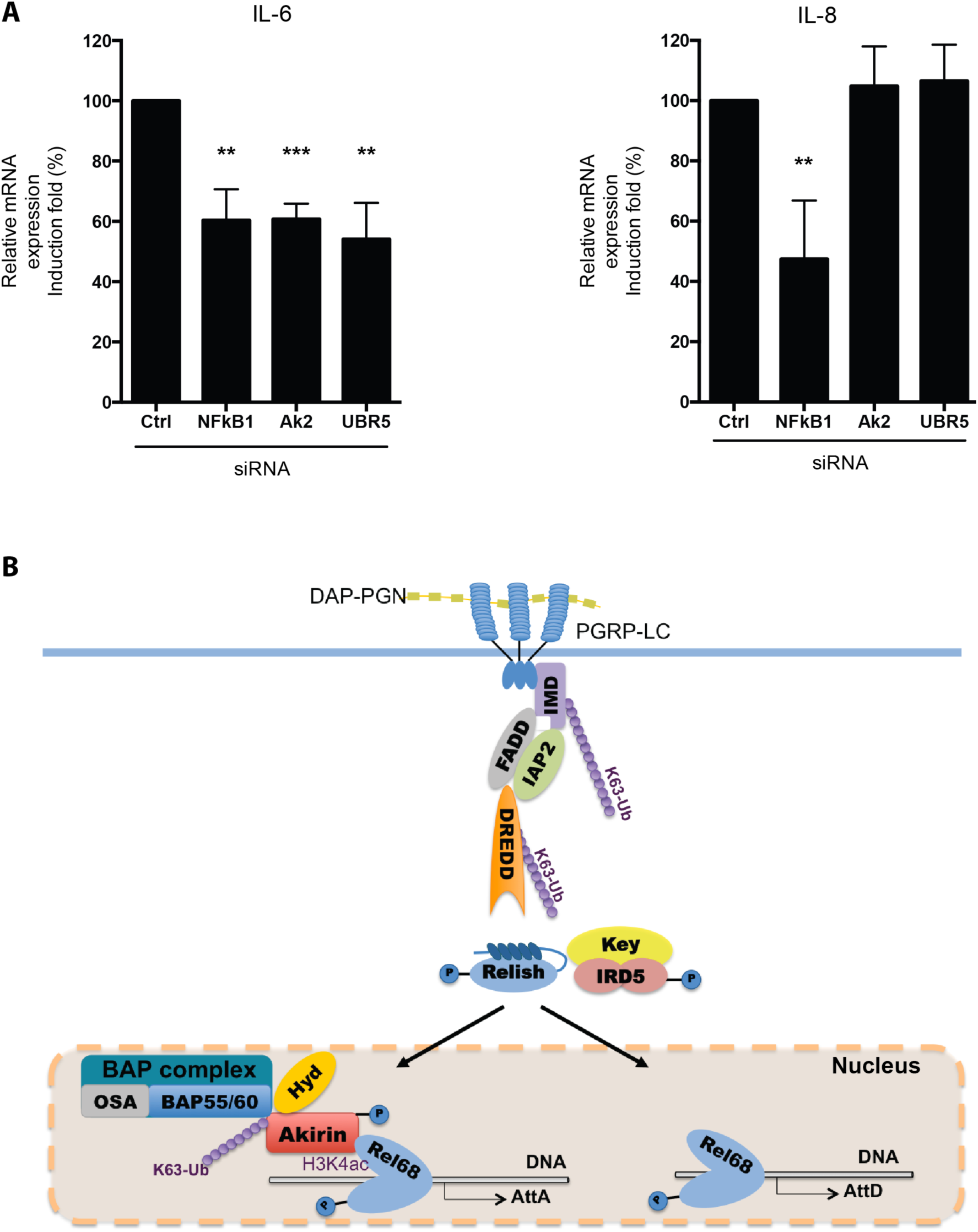
Hyd/Ubr5 is necessary for NF-κB target genes activation. (A) Quantitative RT-PCR of *IL-6* and *IL-8* mRNA from HeLa cells. They were transfected with scrambled siRNA (negative control) or siRNA targeting NFkBl, hAkirin2 (positive controls), and Ubr5. The cells were stimulated with recombinant human IL1β (10 ng/ml) for 4h. Data are represented as mean ± standard deviation of three independent experiments realized on 5 ×10^5^ cells per sample. Statistical significance was established by comparing values from stimulated with unstimulated conditions and genes knockdown with scrambled siRNA control. *P-value < 0.05; **P-value < 0.01; ***P-value < 0.001. Model showing the role of Hyd in the expression of the Akirin-dependent genes in the IMD pathway. After activation of the pathway, allowed by the K63-polyUb chains deposition on the complexes IMD and DREDD by the E3-ubiquitin ligase Iap2, Relish is translocated. The K63-polyUb of Akirin by Hyd allows the protein to link to Relish. This interaction is crucial for the expression of Akirin-dependent genes, necessary for an adequate innate immune response.

Taken altogether, our results show that Hyd/Ubr5 is a HECT E3-ubiquitin ligase involved in NF-κB pathway regulation in *Drosophila* and mammals. In fruit fly, Hyd deposits K63-polyUb chains on Akirin and these ubiquitin marks are required to bridge Akirin and the NF-κB factor Relish. This interaction is necessary for the transcription of an essential NF-κB target genes subset, downstream of the IMD pathway (Fig 4B).

## DISCUSSION

Using *Drosophila* genetics, we describe here for the first time a function for the HECT E3-ubiquitin ligase Hyd in the innate immune response. We could also show using HeLa cells that this function of Hyd downstream of the NF-κB pathway is conserved in humans.

In both humans and *Drosophila*, NF-κB dependent signaling pathways are among the best-known examples of the role of ubiquitin linkage to target proteins in signal transduction (*4, 24*), ubiquitination being involved at every level of the NF-κB pathway, from membrane receptors to chromatin-associated proteins. In order to identify new E3-ubiquitin ligases involved in the *Drosophila* innate immune response, we conducted a RNAi-based screen. We showed, in addition to IAP2 already known to be a bona-fide member of the IMD pathway (*10, 11*), that other RING-domain E3 ubiquitin ligases (CG5334, bon) were involved in the activation of the IMD pathway in *Drosophila* S2 cells after immune challenge. In addition, our results indicate that other RING-domain E3 ubiquitin ligases such as m-cup, Mkrin1 and mura down-regulate IMD pathway target genes activation. Interestingly, this screen also indicates that a HECT E3-ubiquitin ligase, namely Hyd, is involved in the innate immune response.

In *Drosophila*, Hyd was reported to be located in the nuclear and in the cytoplasmic fraction of cells to participate in various phenomenons during development such as cellular proliferation (*17*). More precisely, Hyd shapes hedgehog signaling by differentially restraining the transcriptional activity of Cubitus interuptus via selective association with respective promoters (*23*). And more recently, Hyd and its mammalian orthologue Ubr5 were reported to act at the level of Wnt signaling target genes promoter to enable gene transcription (*25*). Here we identified the HECT E3 ubiquitin-ligase Hyd in *Drosophila* as responsible for the ubiquitination of Akirin and its subsequent binding to the NF-κB transcription factor Relish. Altogether these results point to a conserved function of the HECT E3-ubiquitin ligase Hyd/Ubr5 as a nuclear selector for gene activation.

Downstream of the IMD pathway, the NF-κB transcription factor relish target genes could be divided in two subgroups, Akirin-dependent and Akirin-independent genes (*14*). Targeting Hyd by RNAi in *Drosophila* S2 cells impaired the activation of Akirin-dependent genes upon immune stimulation. Depleting Hyd from *Drosophila* fat-body prevents fly survival to immune challenge with the Gram-negative bacteria *E.coli*, demonstrating the biological relevancy of its function. Co-immunoprecipitation experiments showed that Hyd interacts with Akirin through its catalytic domain to deposit K63-polyUb chains. Of note, the K63-polyubiquitination of Akirin by Hyd is performed only after immune challenge, suggesting that an immune-triggered signal governs this event and remains to be explore.

It is still unclear how the K63-polyubiquitin chains on Akirin physically interact with Relish to set a bridge, as no Ubiquitin Binding Domain (UBD) have been described for Relish. The HECT Ubiquitin ligase family is known in mammals and *Drosophila* to regulate many biological phenomenon (*26*). We found that the mammalian ortholog of Hyd, Ubr5 (*18*) is involved in NF-κB transcriptional selective response in human cell line as well. This suggests a conserved role for Hyd/Ubr5 on hAkirin2, even though we do not know if hAkirin2 is ultimately ubiquitinated. A dedicated study of Ubr5 role in NF-κB pathway is needed to completely assess it. It is known that Ubr5 inhibits the TNF receptor associated factor 3 (Traf3) (*27*) an inhibitor of the NF-κB pathway (*28*). Thus, the role of Ubr5 might be indirect.

When Hyd/Ubr5 is attenuated, only a subset of NF-κB genes is expressed, diminishing the intensity of the innate immune response in *Drosophila* and inflammatory response in mammals, similarly to the inactivation of Akirin (*14, 15*). The link between excessive activation of NF-κB signaling pathway during e.g chronic inflammation and cancer progression or appearance is now on the spotlight (*29*). Uncontrolled activation of NF-κB due to deregulation of ubiquitin-ligases has been reported in many diseases (*30*) and Ubr5 involved in several types of cancer in human (*31*). Our findings point to the HECT E3-ubiquitin ligase Ubr5 as an interesting drug target to modulate NF-κB signaling, control the development of inflammatory diseases and potentially improve treatments of cancer.

## MATERIAL AND METHODS

### Cell culture

S2 cells were cultured at 25°C in Schneider's medium (Biowest) supplemented with 10% fetal calf serum (FCS), penicillin/streptomycin (50 μg/ml of each) and 2 mM glutamax. HeLa cell line was cultured and maintained in DMEM containing 10% (vol/vol) FCS, 40 μg/mL gentamycin. Recombinant human IL1β was purchased from Invitrogen.

### E3-ubiquitin ligases screening methods

A comprehensive list containing 174 E3 ubiquitin ligases in the *Drosophila* genome, consisting predominantly of HECT, RING, and U-box proteins was curated manually by GO- and protein domain-term search in Flybase FB2012_06 Dmel Release 5.48 (*19*). Based on this list, a *Drosophila* E3 ligase dsRNA library was generated in Michael Boutros’s laboratory as previously described (*32*). The screen experiments were performed using 1F3 cells stably expressing AttA firefly luciferase (*12*). Two days after transfection with an Actin renilla luciferase construct, cells were collected and distributed into 96-well screening plates at a density of 4.5 × 10^4^ cells per well. Cells were then transfected with 3 μg of each dsRNA in the *Drosophila* E3-ubiquitin ligase dsRNA library in triplicate by bathing method as previously described (*14*). At day 5 post-transfection, cells were stimulated with heat-killed *E. coli* (40:1) before determining both firefly and renilla luciferase activities.

### RNA interference

The double-strand RNAs for the knockdown experiments in *Drosophila* cells were prepared according to (*14*). Fragments for the different genes were generated from genomic DNA templates using oligonucleotides designed for use with Genome-RNAi libraries (*33*) and are listed in Supplementary Table 1. The small interfering RNAs used for the knockdown experiment in HeLA cells were purchased from Ambion (Supplementary Table S2).

### Luciferase assay

The luciferase assay was realized accordingly to (*14*).

### Plasmid Constructs

pAC-Akirin, pAC-Akirin-V5, pAC-PGRP-LC, pAC-IMD, pMT-Rel-HA and pMT-Bap-HA constructs were described previously (*13, 14*).

### Cell transfection

*Drosophila* S2 cells were transfected with double-strand RNAs using the bathing method described in (*14*) or with plasmids using the Effectene transfection kit (Qiagen). HeLa cells were transfected with siRNA using Lipofectamine RNAiMax (Invitrogen).

### RNA extraction and quantification

For the *ex-vivo* experiments, RNA was extracted from cells and treated with DNAse, using RNA Spin kit (Macherey Nagel). For the *in-vivo* experiments, the procedure was done accordingly to (*14*). Similarly, reverse-transcription and q-RT-PCR were performed as indicated in (*14*). Primers used for q-RT-PCR are listed in Supplementary Table 3.

### Immunoprecipitation and Western blot

The experiments were realized according to (*14*). Immunoprecipitations were performed with rabbit polyclonal anti-Akirin (*14*) and anti-ubiquitin Lys63 specific antibodies (Millipore 05-1308) coupled with Dynabeads Protein G (Invitrogen) and anti-V5 antibodies coupled to agarose beads (Sigma). Proteins were detected by Western blotting using anti-Akirin, anti-ubiquitin Lys63 specific, anti-ubiquitin (Santa cruz biotechnology SC-8017), anti-V5 (Invitrogen r96025), anti-HA (Abcam ab9110) and anti-Relish (gift from Tony Ip) antibodies.

### Fly strains

Stocks were raised on standard cornmeal-yeast-agar medium at 25°C with 60% humidity. To generate conditional knockdown in adult flies, we used the GAL4-GAL80^ts^ system (*22*). Fly lines carrying a UAS-RNAi transgene targeting relish (108469), akirin (109671), and hyd (44675, 44676) were obtained from the Vienna Drosophila RNAi Center (http://stockcenter.vdrc.at/control/main). Fly line carrying a UAS-RNAi transgene against GFP (397-05) was obtained from the Drosophila Genetic Resource Center (Kyoto, Japan; http://www.dgrc.kit.ac.jp/index.html). UAS-RNAi flies were crossed with Actin-GAL4/CyO; Tub-GAL80ts flies at 18°C. Emerged adult flies were then transferred to 29°C to activate the UAS-GAL4 system for 6-7 days.

### Immune challenge

Cells were stimulated with heat-killed *E. coli* (40:1) (*34*). Microbial challenges were performed by pricking adult flies with a sharpened tungsten needle dipped into either PBS or concentrated *Escherichia coli* strain DH5aGFP bacteria solution (*14, 34*). Bacteria were grown in Luria broth (LB) at 29°C.

### Statistical analysis

All P values were calculated using the two-tailed unpaired Student t test (Graph-Pad Prism).

## Acknowledgments

We are grateful to the Drosophila Genomics Resource Center at Indiana University, the Drosophila Genetic Resource Center at the Kyoto Institute of Technology and the Vienna Drosophila RNAi Center for fly stocks.

## Author contributions

N.M., X.-H.N., M.-O.F., A.O. and M.B. designed the experiments. A.C.-M., X.-H.N., A.G. and F.B. performed the experiments. A.C.-M., J.-M.R. and N.M. wrote the manuscript. J.-M.R. and N.M. supervised the study. Competing interests: The authors declare that they have no competing interests.

## SUPPLEMENTARY MATERIALS

**Fig. S1.**
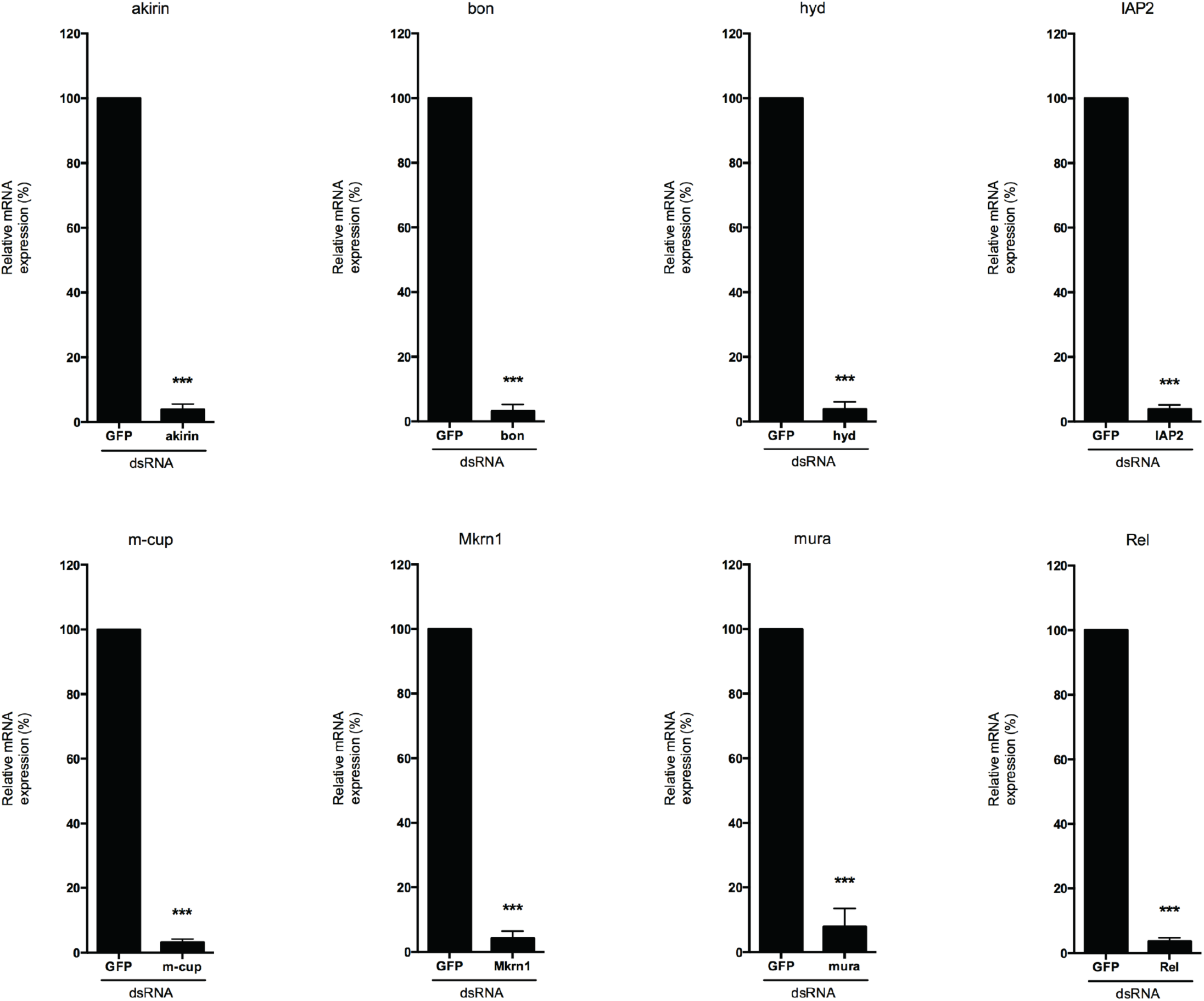
Knockdown efficiency of the double strand RNA used in *Drosophila* S2 cells. Quantitative RT-PCR of akirin, bon, hyd, IAP2, m-cup, Mkm1, mura and relish mRNA from S2 cells transfected with dsRNA against GFP and the respective genes.

**Fig. S2.**
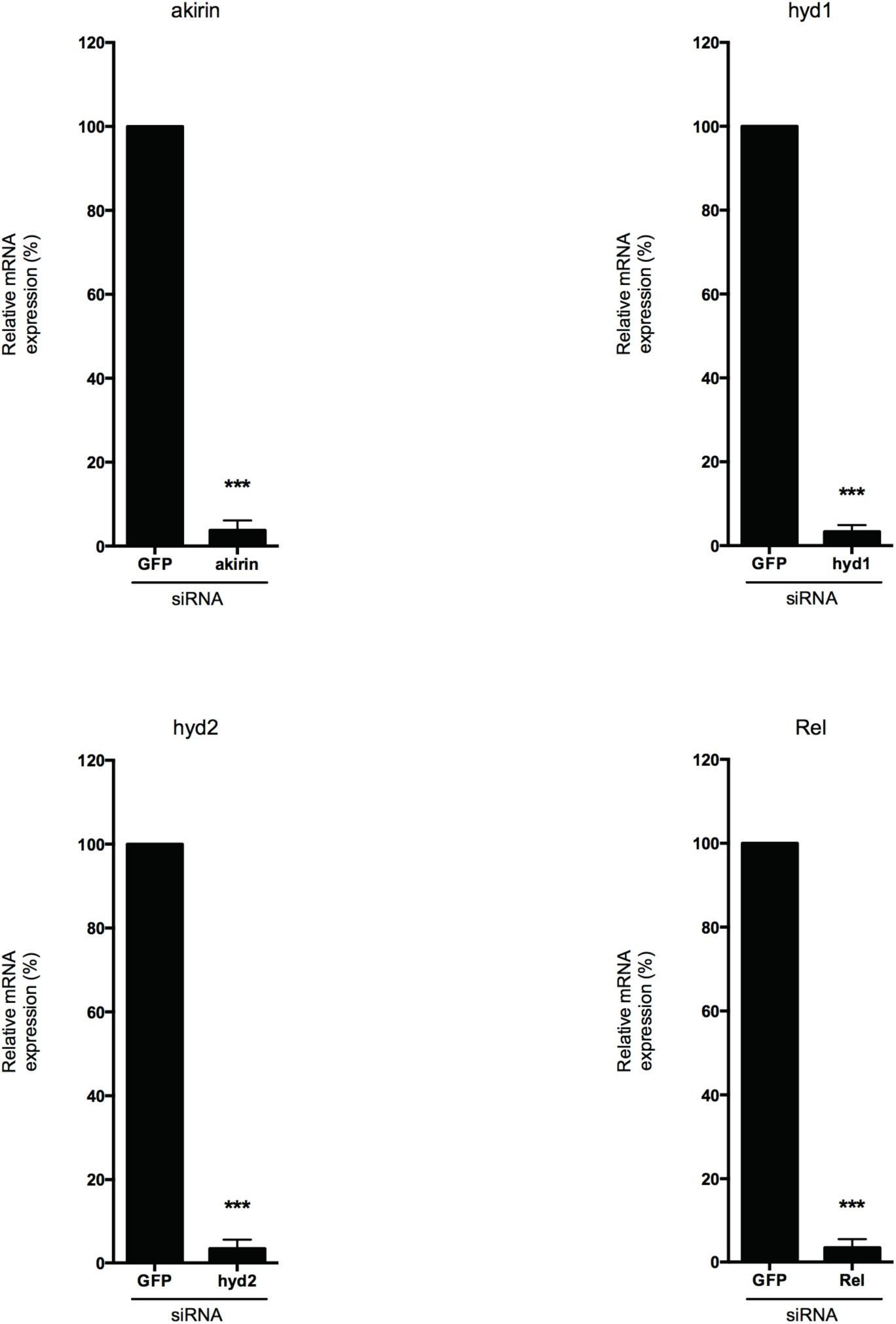
Knockdown efficiency of the Gal4-UAS system used in adult flies. Quantitative RT-PCR of akirin, hyd and relish mRNA from the adult fly lines in which the Gal4-UAS system was used to knockdown the respective genes (two lines for hyd).

**Fig. S3.**
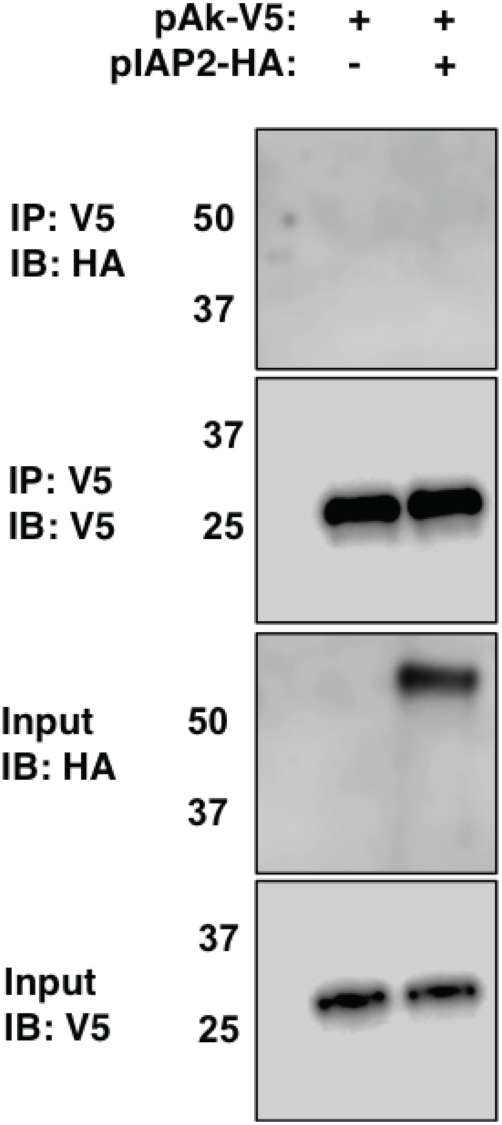
Interaction between IAP2 and Akirin. Co-immunoprecipitation assay between over-expressed IAP2 and Akirin in S2 cells. The cells were transiently transfected with pAC-Akirin-V5 and/or pMT-IAP2-HA. Cell lysates were immunoprecipitated with anti-V5 coupled agarose beads. Immunoprecipitates were analyzed by Western blotting with anti-HA or anti-V5 antibodies. Data are representative of 2 independent experiments.

**Fig. S4.**
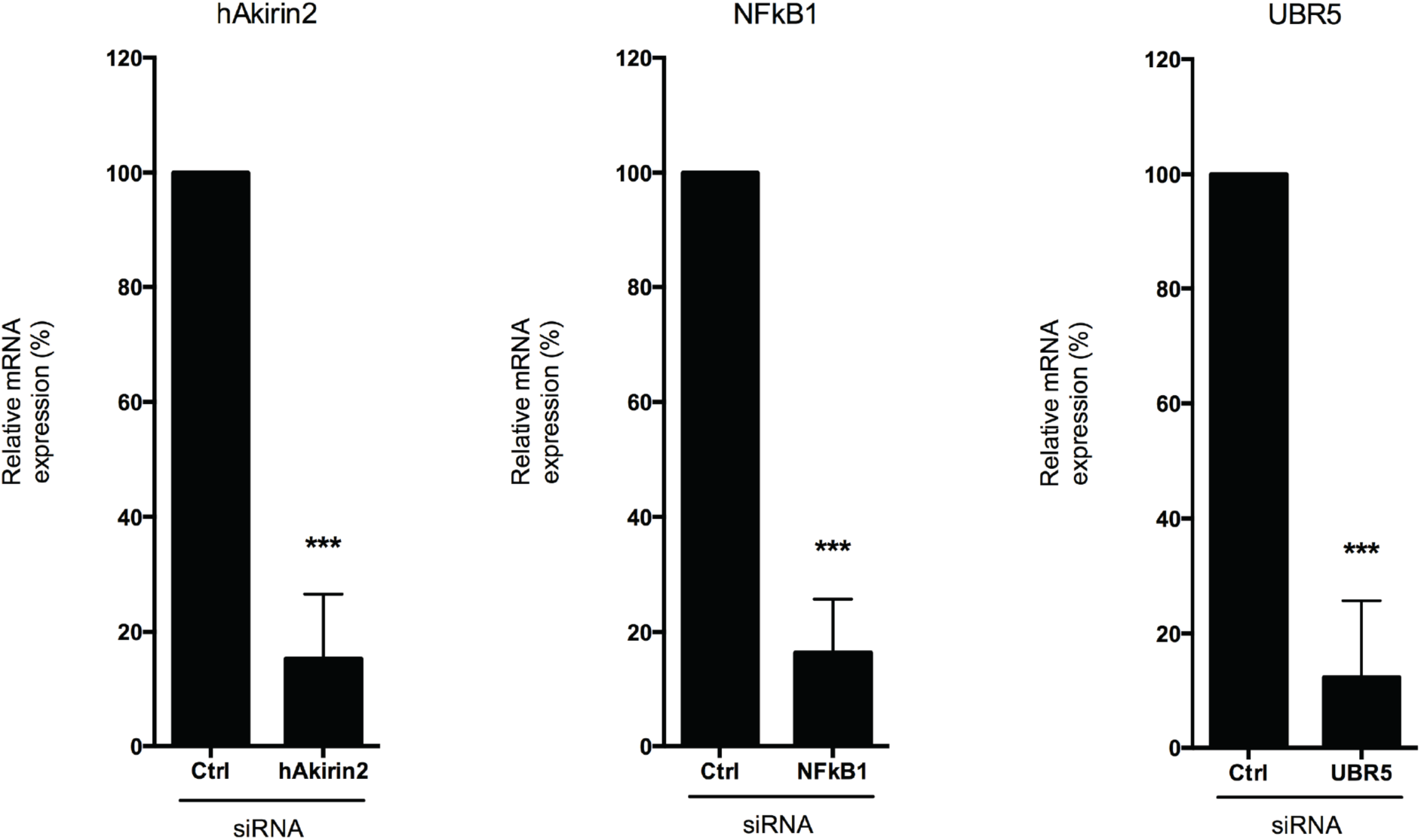
Knockdown efficiency of the small interfering RNA used in HeLa cells. Quantitative RT-PCR of hAkirin2, NFkB1 and Ubr5 mRNA from HeLa cells transfected with scrambled siRNA or targeting the respective genes. Data information: Data for Fig. S1-2 and 6 are represented as mean ± standard deviation of three independent experiments. Statistical significance was established by comparing values from stimulated with unstimulated conditions and genes knockdown with GFP dsRNA or scrambled siRNA control. *P-value < 0.05; **P-value < 0.01; ***P-value < 0.001.

**Table S1.**
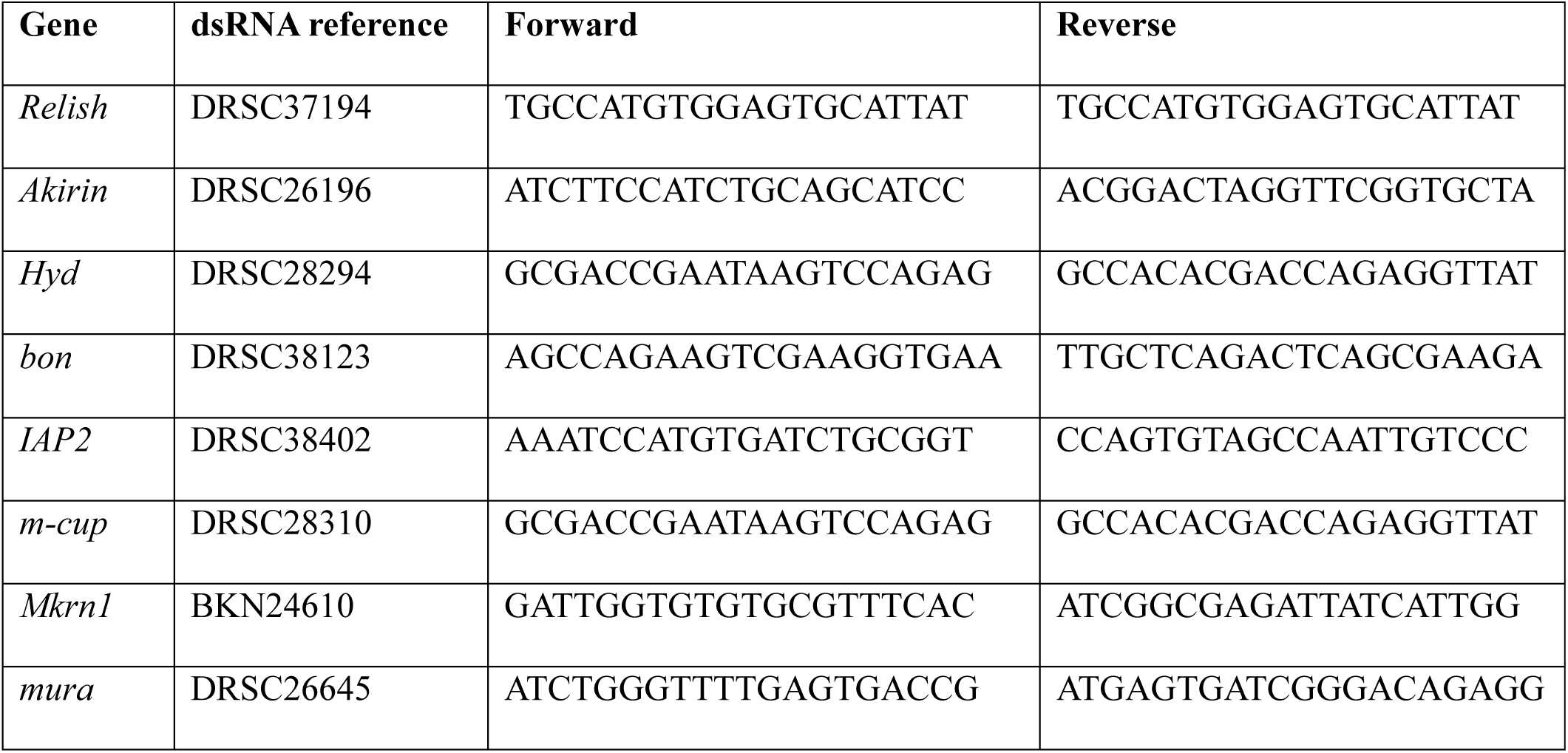
Oligonucleotides used to generate double strand RNA in *Drosophila* S2 cells. Are indicated: gene reference, dsRNA reference (http://www.genomemai.org/GenomeRNAi/), forward and reverse primers (without T7 promoter sequence TTAATACGACTCACTATAGG) used to produce T7 DNA matrix PCR product and PCR product size.

**Table S2.** Oligonucleotides used to generate small interfering RNA in mammalian HeLa cells (Ambion)

| Gene | UniGene ID | siRNA ID |
| --- | --- | --- |
| *Negative Control* | - | AM4611 |
| *NFkB1* | Hs.618430 | s9504 |
| *Ak2* | Hs.485915 | s30221 |
| *UBR5* | Hs.492445 | s224201 |

**Table S3.**
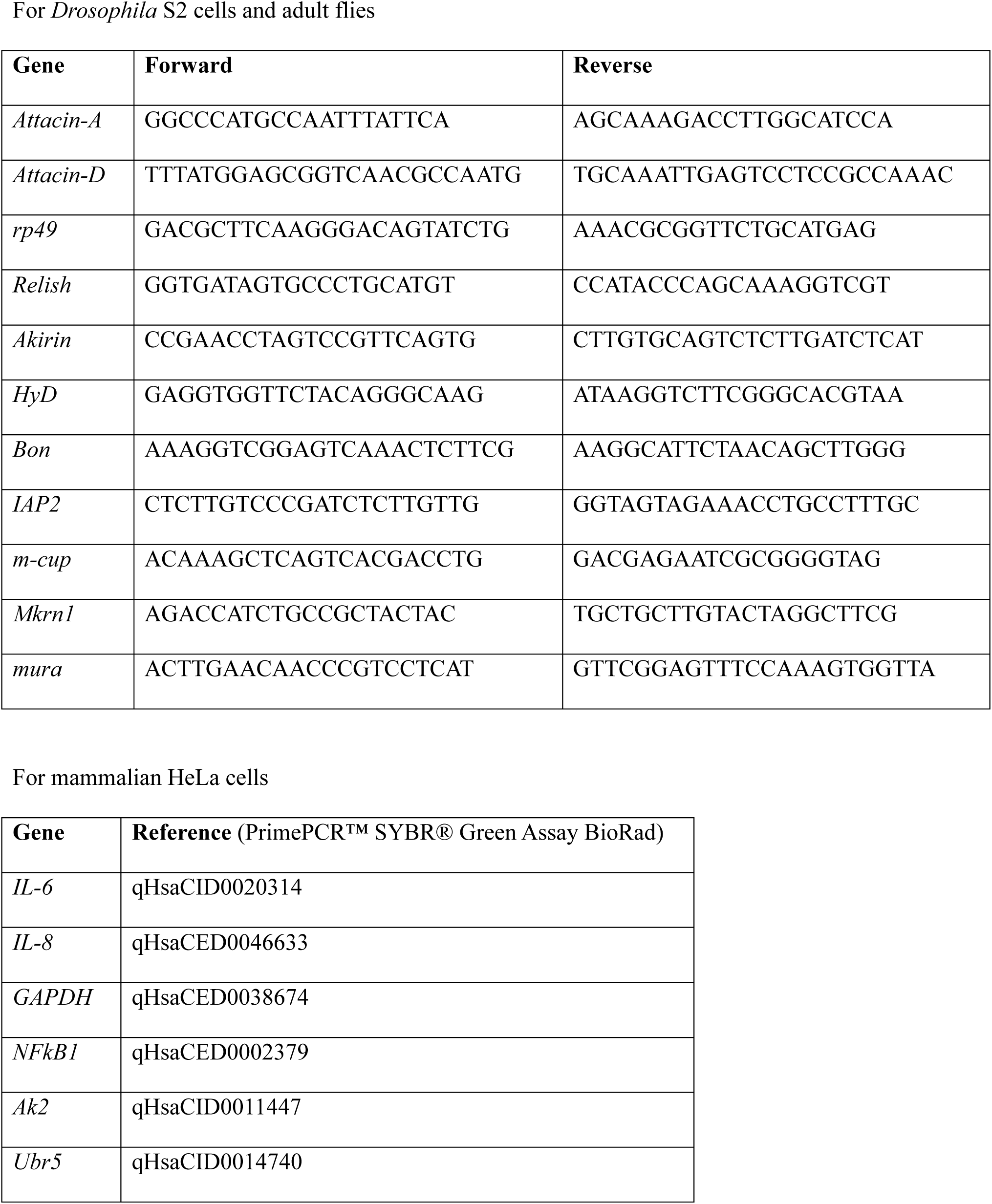
Oligonucleotides used for quantitative real-time PCR.

## REFERENCES AND NOTES

1. M. Vidal, R. L. Cagan, Drosophila models for cancer research. Current opinion in genetics & development 16, 10–16 (2006).

2. S. Maeda, M. Omata, Inflammation and cancer: role of nuclear factor-kappaB activation. Cancer Sci 99, 836–842 (2008).

3. D. Ferrandon, J. L. Imler, C. Hetru, J. A. Hoffmann, The Drosophila systemic immune response: sensing and signalling during bacterial and fungal infections. Nature reviews. Immunology 7, 862–874 (2007).

4. Y. Park, H. S. Jin, D. Aki, J. Lee, Y. C. Liu, The ubiquitin system in immune regulation. Advances in immunology 124, 17–66 (2014).

5. K. N. Swatek, D. Komander, Ubiquitin modifications. Cell research 26, 399–422 (2016).

6. D. Thevenon et al., The Drosophila ubiquitin-specific protease dUSP36/Scny targets IMD to prevent constitutive immune signaling. Cell host & microbe 6, 309–320 (2009).

7. I. Aksentijevich, Q. Zhou, NF-kappaB Pathway in Autoinflammatory Diseases: Dysregulation of Protein Modifications by Ubiquitin Defines a New Category of Autoinflammatory Diseases. Frontiers in immunology 8, 399 (2017).

8. M. G. Kattah, B. A. Malynn, A. Ma, Ubiquitin-Modifying Enzymes and Regulation of the Inflammasome. Journal of molecular biology 429, 3471–3485 (2017).

9. L. H. Gallo, J. Ko, D. J. Donoghue, The importance of regulatory ubiquitination in cancer and metastasis. Cell Cycle 16, 634–648 (2017).

10. A. Kleino et al., Inhibitor of apoptosis 2 and TAK1-binding protein are components of the Drosophila Imd pathway. EMBO J 24, 3423–3434 (2005).

11. V. Gesellchen, D. Kuttenkeuler, M. Steckel, N. Pelte, M. Boutros, An RNA interference screen identifies Inhibitor of Apoptosis Protein 2 as a regulator of innate immune signalling in Drosophila. EMBO Rep 6, 979–984 (2005).

12. H. Fukuyama et al., Landscape of protein-protein interactions in Drosophila immune deficiency signaling during bacterial challenge. Proc Natl Acad Sci U S A 110, 10717–10722 (2013).

13. A. Goto et al., Akirins are highly conserved nuclear proteins required for NF-kappaB-dependent gene expression in drosophila and mice. Nature immunology 9, 97–104 (2008).

14. F. Bonnay et al., Akirin specifies NF-kappaB selectivity of Drosophila innate immune response via chromatin remodeling. EMBO J, (2014).

15. S. Tartey et al., Akirin2 is critical for inducing inflammatory genes by bridging IkappaB-zeta and the SWI/SNF complex. EMBO J, (2014).

16. J. D. Lee, K. Amanai, A. Shearn, J. E. Treisman, The ubiquitin ligase Hyperplastic discs negatively regulates hedgehog and decapentaplegic expression by independent mechanisms. Development 129, 5697–5706 (2002).

17. E. Mansfield, E. Hersperger, J. Biggs, A. Shearn, Genetic and molecular analysis of hyperplastic discs, a gene whose product is required for regulation of cell proliferation in Drosophila melanogaster imaginal discs and germ cells. Developmental biology 165, 507–526 (1994).

18. M. J. Callaghan et al., Identification of a human HECT family protein with homology to the Drosophila tumor suppressor gene hyperplastic discs. Oncogene 17, 3479–3491 (1998).

19. L. S. Gramates et al., FlyBase at 25: looking to the future. Nucleic Acids Res 45, D663–D671 (2017).

20. S. Tauszig, E. Jouanguy, J. A. Hoffmann, J. L. Imler, Toll-related receptors and the control of antimicrobial peptide expression in Drosophila. Proc Natl Acad Sci U S A 97, 10520–10525 (2000).

21. A. Kleino, N. Silverman, The Drosophila IMD pathway in the activation of the humoral immune response. Developmental and comparative immunology 42, 25–35 (2014).

22. S. E. McGuire, G. Roman, R. L. Davis, Gene expression systems in Drosophila: a synthesis of time and space. Trends Genet 20, 384–391 (2004).

23. G. Wang et al., Hyperplastic discs differentially regulates the transcriptional outputs of hedgehog signaling. Mech Dev 133, 117–125 (2014).

24. S. Ghosh, J. F. Dass, Study of pathway cross-talk interactions with NF-kappaB leading to its activation via ubiquitination or phosphorylation: A brief review. Gene 584, 97–109 (2016).

25. J. E. Flack, J. Mieszczanek, N. Novcic, M. Bienz, Wnt-Dependent Inactivation of the Groucho/TLE Co-repressor by the HECT E3 Ubiquitin Ligase Hyd/UBR5. Molecular cell 67, 181–193 e185 (2017).

26. M. Scheffner, S. Kumar, Mammalian HECT ubiquitin-protein ligases: biological and pathophysiological aspects. Biochimica et biophysica acta 1843, 61–74 (2014).

27. J. H. Cho et al., The p90 ribosomal S6 kinase-UBR5 pathway controls Toll-like receptor signaling via miRNA-induced translational inhibition of tumor necrosis factor receptor-associated factor 3. J Biol Chem 292, 11804–11814 (2017).

28. J. Q. He, S. K. Saha, J. R. Kang, B. Zarnegar, G. Cheng, Specificity of TRAF3 in its negative regulation of the noncanonical NF-kappa B pathway. J Biol Chem 282, 3688–3694 (2007).

29. K. Taniguchi, M. Karin, NF-kappaB, inflammation, immunity and cancer: coming of age. Nature reviews. Immunology 18, 309–324 (2018).

30. K. Iwai, Diverse roles of the ubiquitin system in NF-kappaB activation. Biochimica et biophysica acta 1843, 129–136 (2014).

31. R. F. Shearer, M. Iconomou, C. K. Watts, D. N. Saunders, Functional Roles of the E3 Ubiquitin Ligase UBR5 in Cancer. Mol Cancer Res 13, 1523–1532 (2015).

32. M. Boutros et al., Genome-wide RNAi analysis of growth and viability in Drosophila cells. Science 303, 832–835 (2004).

33. E. E. Schmidt et al., GenomeRNAi: a database for cell-based and in vivo RNAi phenotypes, 2013 update. Nucleic Acids Res 41, D1021–1026 (2013).

34. J. M. Reichhart, D. Gubb, V. Leclerc, The Drosophila serpins: multiple functions in immunity and morphogenesis. Methods in enzymology 499, 205–225 (2011).

